# Impact and centrality of scientific disciplines

**DOI:** 10.1101/2023.09.01.555991

**Authors:** Daniel Aguilar-Velázquez, Rodolfo Romero Herrera, Denis Boyer, Gabriel Ramos-Fernández

## Abstract

The Scimago Journal Rank (SJR) is a metric that captures the centrality of a journal across an all-discipline article network, while the impact factor (IF) is the average incoming citations of a journal. We analyzed SJRs and IFs of the journals belonging to the SJR first quartile from 2013 to 2020 in 7 disciplines: mathematics, biology, physics, medicine, social sciences, chemistry, and engineering. We show that biology is the most central discipline, followed by physics and chemistry. These three disciplines also present the highest IFs. Mathematics journals display a low IF (the second-lowest among disciplines), but possesses an intermediate centrality. While the average IF has increased over the last years, the SJR average has decreased. Gini coefficients show that SJR is a slightly more egalitarian metric than IF. We discuss some possible origins of these findings.

## Introduction

The definition of the impact factor is clear and intuitive: the number of incoming citations to a journal divided by the number of papers published (see methods). IF and other citations metrics are used by databases (Scopus and Web of Science) to classify scientific journals. In addition, IF is used by employers to evaluate and hire scientists. The IF has taken a controversial role in the evaluation of scientific journals because there are great differences across disciplines [1]. For example, a top journal in mathematics may show an impact factor of 2, whereas top medicine and biology journals can reach values larger than 50.

A metric that complements the impact factor is the Scimago Journal Rank. SJRs are obtained from applying the PageRank algorithm to the Journal citation network [2, 3]. This algorithm measures the centrality of a journal taking into account the importance/impact of the incoming citation. Therefore, we can observe journals of high centrality and low IF. Using principal component analysis some authors have shown that journal metrics are principally distributed in two clusters: one cluster is related to the number of citations per document and the other cluster comprises PageRank and centrality measures [4]. Moreover, betweenness centrality and PageRank have been linked to interdisciplinarity [5, 6]. IF and SJR depend on the graph properties of the citation network. The equivalent representation of the impact factor in a citation network is the instrength, which depends on the number of citations, whereas SJR depends on the centrality and impact of the incoming citations (Fig. 1). In this study, we compare impact and centrality patterns among scientific disciplines.

**Fig 1.**
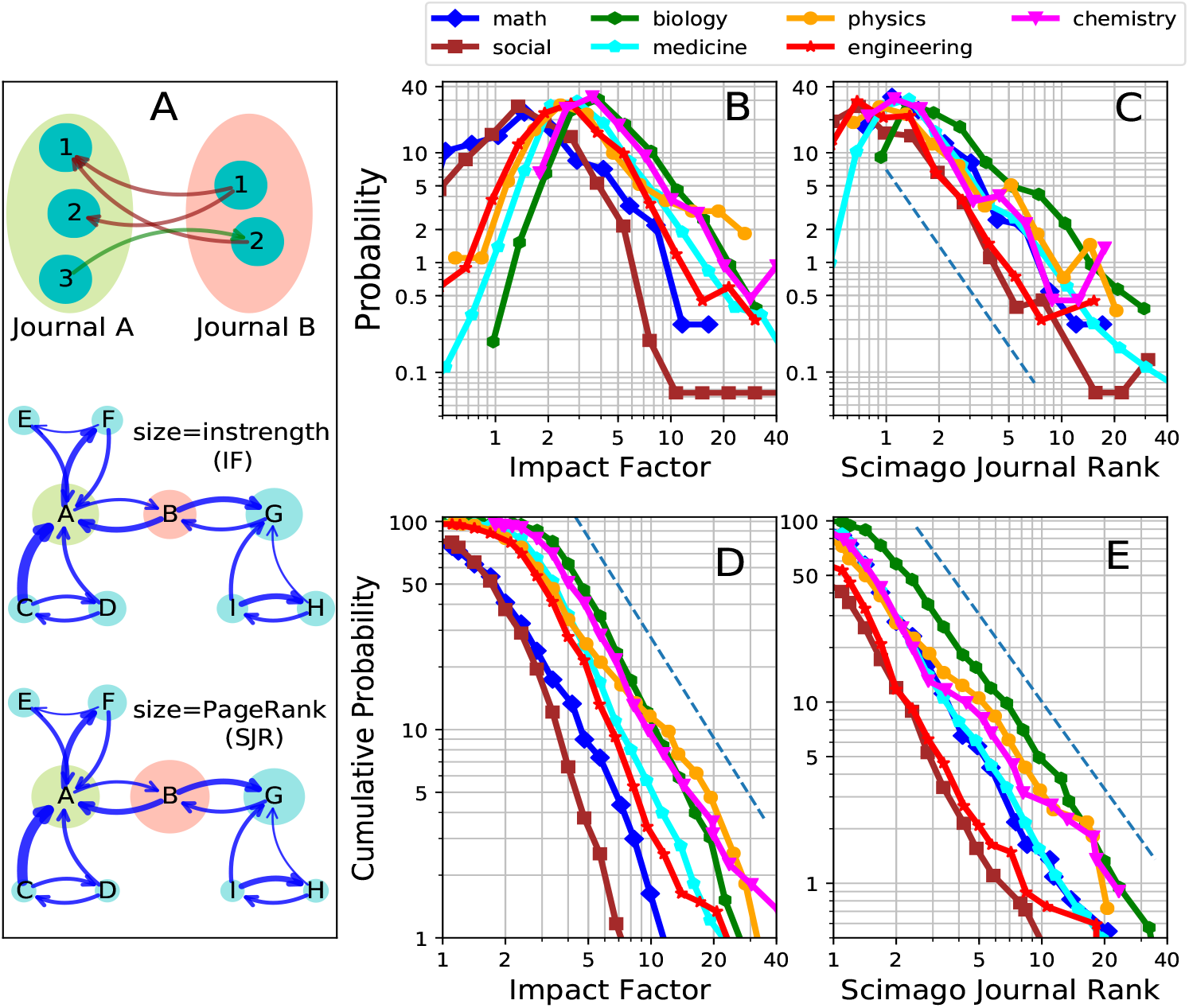
IF and SJR distributions. A) Schematic representation of IF represented by the instrength and SJR represented by PageRank. Top: nodes are articles and links are citations. Middle and bottom: nodes are journals and links are citations. B) IF probability distribution. C) SJR probability distribution. The dashed line indicates a power-law function *P* (*SJR*) = *SJR*^*−*2.4^. D) IF and E) SJR cumulative probability (CP). The dashed line indicates a power-law function *CP* (*SJR*) = *SJR*^*−*1.6^.

## Methods

The IF is the average number of citations to articles published in the previous two years. IF is computed as follows:

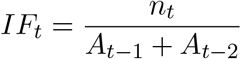

where *A*_*t−*1_ and *A*_*t−*2_ are the number of articles published during year t-1 and t-2, respectively. *n*_*t*_ is the number of citations these articles received at year *t*.

The SJR metric is based on the PageRank algorithm [2], which measures the centrality of a given node in a network. It is computed through the following equation [7]:

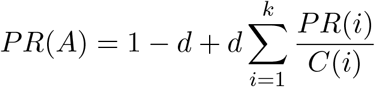

Where *PR*(*A*) is the PageRank of the node *A, d* is the damping factor that is commonly assigned a value of 0.85, *k* is the indegree, *PR*(*i*) is the PageRank of the incoming links (citations), and *C*(*i*) is the number of outgoing links (references). In the algorithm, all the journals are initialized with the same PageRank value and then one proceeds to update recursively. PageRank is strongly related to random walk centrality [8].

We computed the Gini coefficient using the formula [9],

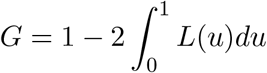

where *L*(*u*) is the Lorentz curve that represents the cumulative probability. A Gini coefficient is equal to 0 when all journals posses either the same IF or SJR value. One obtains a Gini coefficient close to 1 when the total IF or SJR is concentrated in a few journals.

## Results

We extracted data from SJR databases. We analyze 5184 SJRs and IFs for 2013, 5414 for 2017, and 5796 for 2020. Scimago also computes the average citation per document in a 2-year period, i.e. the IF [10]. Fig. 1A (top) shows citations among articles belonging to two journals, where the number of citations determined the strength from journal A to journal B and vice versa (middle), and where the instrength represents the IF. In Fig. 1A (bottom), journal B possesses few citations but a high SJR. Fig. 1B shows the probability distributions of the IFs for seven disciplines for the 2017 year. We observe that the most probable value for mathematics and social science is around 1.5, whereas it is close to 3.0 for physics, medicine and engineering, and 4.0 for biology and chemistry. In Fig 1C, the tails of the distributions approximately follow a power-law function *P* (*SJR*) = *SJR*^*−*2.4^ for SJR¿ 2. Next, we display the cumulative probability (CP), or the probability that IF ¿ value (Fig. 1D). Biology, chemistry, and physics have the “fattest” tails with a significant amount of high IF journals. For example, more than 80% of biology journals score an IF¿3.0, whereas only 20% of math journals meet this condition. Social sciences show the lowest probability that IF¿3. The cumulative probabilities of SJR follow a power-law behavior *CP* (*SJR*) = *SJR*^*−*1.6^ (Fig. 1E). Biology is the most central field, with the highest SJR values, followed by physics and chemistry, similarly to their IF patterns. In contrast, maths are more central than engineering and as central as medicine, despite having a lower mean IF, see Fig. 1D.

The evolution of the IFs and SJRs from 2013 to 2020 is studied in Fig. 2. The scatter plot of Fig. 2A displays the IF changes of 3927 journals between 2013 and 2020, resulting in an overall increase with a rate of 4.51% per year during this period. For the SJRs, an overall decrease of 2.13% per year is found over the 2017 to 2020 period (Fig. 2B). From Fig. 2C, one notices that all the seven disciplines increased their average impact between 2013 and 2017, whereas only medicine decreased between 2017 and 2020. Conversely, the average SJR decreased (Fig. 2D) between 2017 and 2020 in all disciplines except engineering (it remained constant for medicine). The SJR and IF averaged over all disciplines (Fig. 2E) reveal a steady growth of the impact from 2013 and a drop in centrality, mostly in recent years. Fig. 2F shows the evolution of the Gini coefficient, which gives information on the inequality of a distribution, corresponding to the SJRs and IFs of all journals (solid lines) or averaged over the different disciplines. We found that in general, SJR Gini coefficients are substantially smaller than IF Gini coefficients, indicating that SJRs are more equally distributed than IFs. In addition, averages over all disciplines reveal that the IF has become more unequal, whereas SJR is becoming more equally distributed.

**Fig 2.**
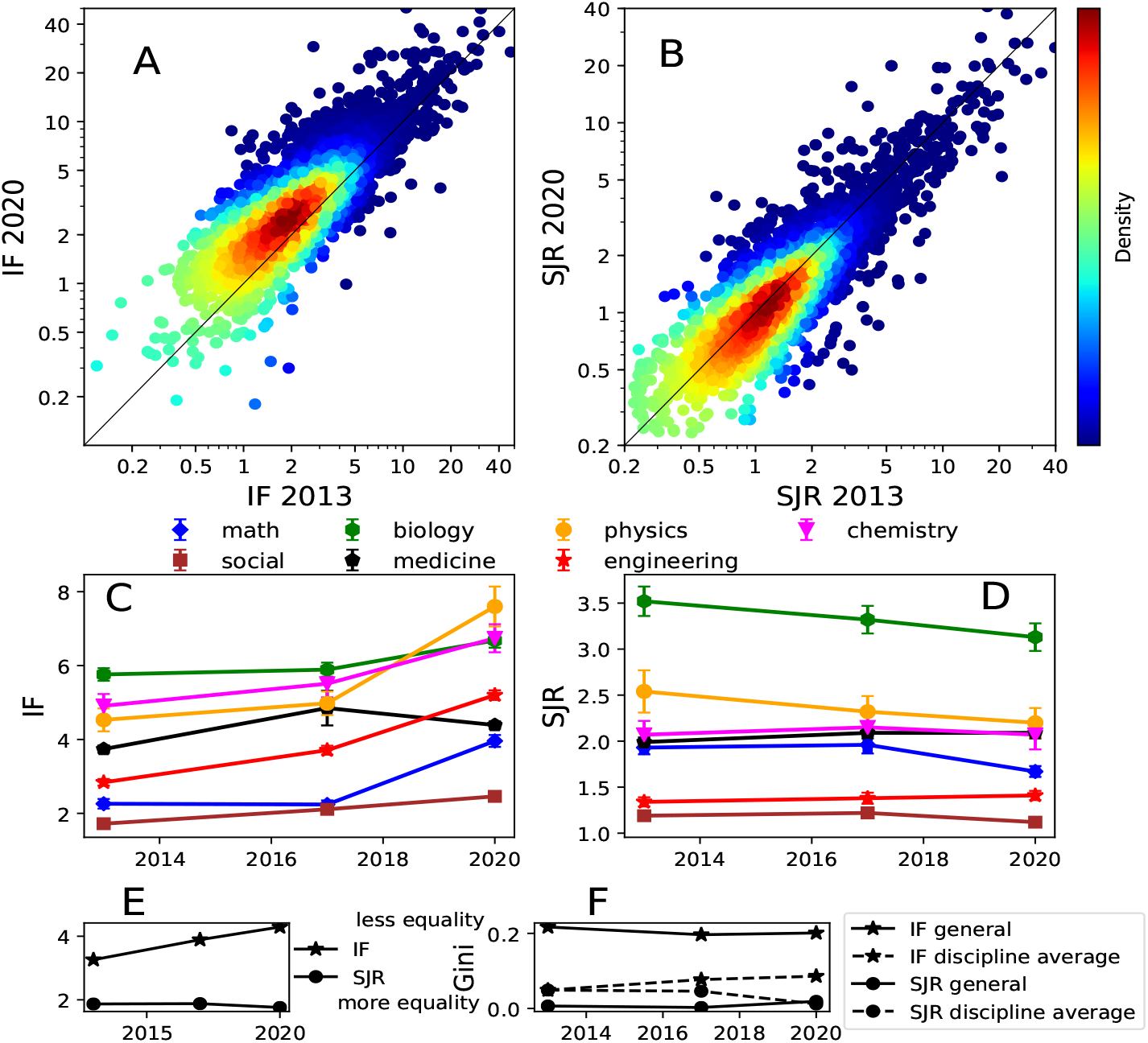
Changes in IF and SJR from 2013 to 2020. A) Density scatter plot of 2013 IF vs 2020 IF for 3927 journals belonging to Q1 from the seven disciplines. B) Same as A) for SJR. Average IF (C) and SJR (D) for the seven disciplines. Error bars represent mean *±* standard error of the mean. E) Average IF and SJR across all disciplines in the 2013-2020 period. F) Gini coefficient for IF and SJR.

## Discussion

Previous findings indicate that biology exhibits a high interdisciplinarity, as measured by the number of references made and citations received from other disciplines [11]. This is consistent with our result that biology is the most central discipline. IF is dominated by biology, physics and chemistry. Although the average IF has increased from 2013 to 2020 (following the trend observed between 1994 and 2005, see [1]), the average SJR has decreased from 2017 to 2020. The IF increment was attributed to an increase of the average number of references [1]. Concerning the SJR, PageRank represents the probability to cite a given journal [7], and the Q1 SJR decrease may be caused by the SJR increment of other journals, either belonging to other disciplines or quartiles. For example, Q3 and Q4 SJRs increased from 2013 to 2020 (see suplementary information). It is important to mention some limitations of our study, such as the lack of access to the journal network from which SJRs and IFs are obtained. In addition, the number of references in articles varies across disciplines and may influence IFs and SJRs. We also find that mathematics show a high centrality compared to its impact and an opposite effect for engineering. Although each discipline contains citations among the same discipline, the results of the IF may be partially explained by an implicit sequence: maths-physics-chemistry-biology. Namely, maths solves problems in physics, chemistry, and biology. Physics explains some phenomena in chemistry, and chemistry governs many biological processes. For example, Erwin Schroödinger anticipated the DNA structure in his “What is life” lecture series [12], the mathematician Alan Turing worked on artificial intelligence and morphogenesis, producing seminal contributions on these text topics [13, 14]. This tendency remains to recent publications [15–17]. In this way, we expect that biology receives the largest number of citations, followed by chemistry, physics and maths. Our explanatory sequence is supported by the IF evolution between 2013 and 2017 (fig. 2C), however in 2020, physics displays the highest IF. In contrast, SJR is less related to the above explanatory sequence, as physics is more central than chemistry. Some maths and physics breakthrough contributions, like Watts and Strogatz ‘s article introducing the small-world concept, received thousands of citations from multiple disciplines [18]. IF variations could be related to the differences between abstract (maths) and empirical (biology) knowledge: it is more difficult to explain to an audience a new mathematical advance than a biological or medicinal discovery. This aspect could contribute to the differences observed among disciplines in journal metrics.

## Supporting information

Supplementary Information for Impact and centrality of scientific disciplines

## Data availability

Data is available at Scimago Journal Rank web page: https://www.scimagojr.com/journalrank.php

## Acknowledgments

This work was partially supported by Consejo Nacional de Ciencia y Tecnología (CONACYT).

## Notes

### Competing Interest Statement

The authors have declared no competing interest.

