## Supplementary Information for Impact and centrality of scientific disciplines for "Impact and centrality of scientific disciplines"

This PDF file includes:

Supplementary text

Supplementary figure

The evolution of the Q3 and Q4 SJRs from 2013 to 2020 is shown at Fig. S1 (top). The scatter plot comprises changes of 6515 journals between 2013 and 2020. The Q2, Q3, and Q4 SJR averaged over all disciplines is shown in Fig. S1 (bottom). For Q4, SJRs increase with a rate of 1.03% per year during 2013-2017 period, and 3.76% for 2017-2020 period. Q3 SJRs increase with a rate of 1.9% per year during 2013-2017 period, and 4.3% for 2017-2020 period. Q2 SJRs register a increase of 1.12% during 2013-2017 period, and remain constant for 2017-2020 period.

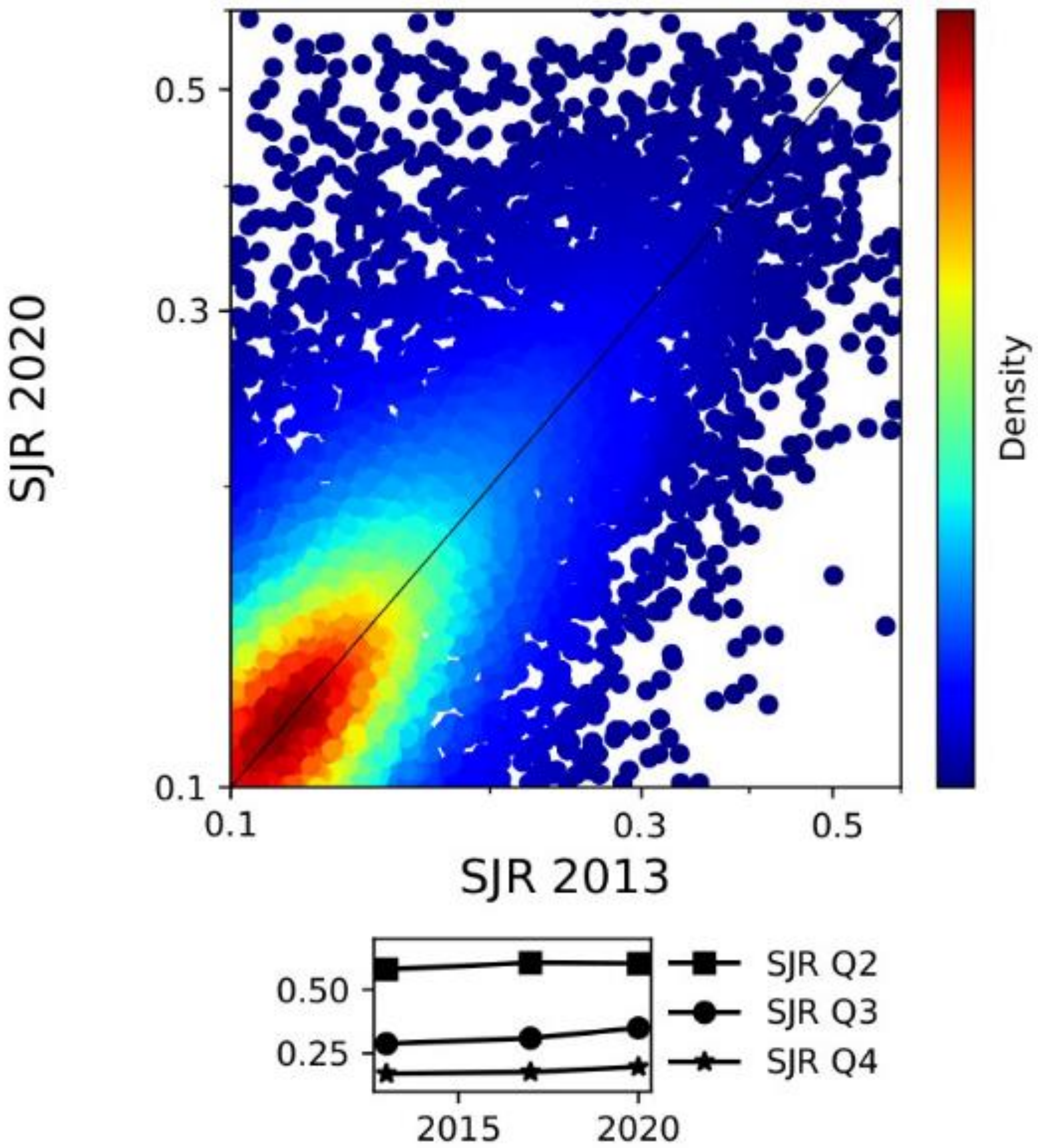


Fig. S1. Changes in Q2, Q3 and Q4 SJRs from 2013 to 2020. Top: density scatter plot of 2013 SJR vs 2020 SJR for 6515 journals belonging to Q3 and Q4 from the seven disciplines. Average SJR across all disciplines in the 2013-2020 period.
